# Temporal-spectral signaling of sensory information and expectations in the cerebral processing of pain

**DOI:** 10.1101/2021.06.24.449790

**Authors:** Moritz M. Nickel, Laura Tiemann, Vanessa D. Hohn, Elisabeth S. May, Cristina Gil Ávila, Falk Eippert, Markus Ploner

**Affiliations:** Department of Neurology, School of Medicine, Technical University of Munich (TUM), Munich, Germany; TUM-Neuroimaging Center, School of Medicine, TUM, Munich, Germany; Max Planck Institute for Human Cognitive and Brain Sciences, Leipzig, Germany; Center for Interdisciplinary Pain Medicine, School of Medicine, TUM, Munich, Germany

## Abstract

The perception of pain is shaped by somatosensory information about threat. However, pain is also influenced by an individual’s expectations. Such expectations can result in clinically relevant modulations and abnormalities of pain. In the brain, sensory information, expectations (predictions), and discrepancies thereof (prediction errors) are signaled by an extended network of brain areas. These brain areas generate evoked potentials and oscillatory responses at different latencies and frequencies. Recent evidence has provided first insights into how oscillatory responses at different frequencies signal predictions and prediction errors. However, a comprehensive picture of how evoked and oscillatory brain responses signal sensory information, predictions, and prediction errors in the processing of pain is lacking so far. We therefore built upon a neuroimaging study which investigated the spatial signalling of sensory information, predictions and predictions errors in the processing of pain (Geuter et al., 2017). To complement and extend this study, we applied brief painful stimuli to 48 healthy human participants and independently modulated sensory information (stimulus intensity) and expectations of pain intensity while measuring brain activity using electroencephalography (EEG). Pain ratings confirmed that pain intensity was shaped by both sensory information and expectations. In contrast, Bayesian analyses revealed that stimulus-induced EEG responses at different latencies (the N1, N2, and P2 components) and frequencies (alpha, beta, and gamma oscillations) were shaped by sensory information but not by expectations. Expectations, however, shaped alpha and beta oscillations before the painful stimuli. These findings indicate that commonly analyzed EEG responses to painful stimuli are more involved in signaling sensory information than in signaling expectations or mismatches of sensory information and expectations. Moreover, they indicate that the effects of expectations on pain are served by brain mechanisms which differ from those conveying effects of sensory information on pain.

## Introduction

The perception of pain emerges from the integration of sensory information about threat and contextual factors such as an individual’s expectations (Atlas and Wager, 2012; Peerdeman et al., 2016a; Fields, 2018). For instance, expectations of pain relief during placebo manipulations can yield substantial and clinically highly relevant decreases of pain (Finniss et al., 2010; Enck et al., 2013; Wager and Atlas, 2015). Moreover, expectations cannot only alleviate pain but also significantly influence the development (Vlaeyen et al., 2016) and prognosis of chronic pain (Cormier et al., 2016; Peerdeman et al., 2016b). Thus, understanding how the brain translates sensory information and expectations into pain promises important insights into the neural mechanisms of pain in health and disease.

In the brain, pain is associated with the activation of an extended network of brain areas (Garcia-Larrea and Peyron, 2013; Baliki and Apkarian, 2015) which yields electrophysiological responses at different latencies and frequencies (Ploner and May, 2018). These responses comprise evoked potentials including the early N1 and later N2 and P2 components (Garcia-Larrea et al., 2003; Lorenz and Garcia-Larrea, 2003) as well as oscillatory responses at alpha (8-13 Hz), beta (13-30Hz) and gamma (40-100 Hz) frequencies (Ploner et al., 2017). Electroencephalography (EEG) and magnetoencephalography (MEG) studies have provided important insights into the functional significance of these responses. The early N1 response has been particularly related to objective sensory information while the later N2 and P2 components (Garcia-Larrea et al., 1997; Lee et al., 2009) as well as gamma oscillations (Gross et al., 2007; Hu and Iannetti, 2019) are more closely related to subjective pain perception. However, results on how expectations shape the different responses are inconsistent (Lorenz et al., 2005; Wager et al., 2006; Brown et al., 2008; Colloca et al., 2008; Iannetti et al., 2008; Morton et al., 2010; Lyby et al., 2011; Huneke et al., 2013; Tiemann et al., 2015; Hird et al., 2018). Thus, a comprehensive assessment of how different evoked and oscillatory brain responses signal sensory information, expectations, and pain is lacking so far.

Moving beyond the domain of pain, the predictive coding framework of brain function (Huang and Rao, 2011; Clark, 2013) is a general theory used to describe the encoding and integration of sensory information and expectations (de Lange et al., 2018). The predictive coding framework proposes that the brain continuously generates predictions about the environment. These predictions are compared against sensory evidence and discrepancies produce prediction errors that serve to optimize future predictions. In this way, the brain efficiently allocates its limited resources to events that are behaviorally relevant and useful for updating predictions (i.e., learning processes) (Friston, 2010). It has been suggested that predictive coding processes are implemented by evoked potentials at different latencies (Friston, 2005) and neuronal oscillations at different frequencies (Arnal and Giraud, 2012; Bastos et al., 2012). In particular, it has been shown that already the earliest evoked potential components are shaped by predictions (Rauss et al., 2011; Bendixen et al., 2012), whereas later responses have been related to prediction errors (Stefanics et al., 2018). Moreover, alpha and beta oscillations have been implicated in the signaling of predictions, whereas gamma oscillations have been proposed to signal prediction errors (Arnal and Giraud, 2012; Bastos et al., 2012; Bastos et al., 2015; Michalareas et al., 2016; Sedley et al., 2016).

Considering the preeminent role of the integration of sensory information and expectations in the processing of pain, an application of predictive coding frameworks to pain is obvious (Buchel et al., 2014; Ploner et al., 2017; Tabor et al., 2017; Ongaro and Kaptchuk, 2019; Seymour, 2019). This is even more appealing as abnormally precise predictions and/or abnormal updating of predictions might figure prominently in the pathology of chronic pain (Edwards et al., 2012; Wiech, 2016; Henningsen et al., 2018). Consequently, recent fMRI studies have applied predictive coding frameworks to the processing of pain (Geuter et al., 2017; Fazeli and Buchel, 2018). The results revealed a spatial dissociation of stimulus intensity coding and predictions coding. For instance, in the insular cortex, a striking posterior-to-anterior gradient from the encoding of stimulus intensity to the encoding of predictions and prediction errors was observed. Most recently, a first EEG study applied a predictive coding framework to oscillatory responses to noxious stimuli (Strube et al., 2021). The findings indicated that alpha-to-beta and gamma oscillations signal expectations and prediction errors in the processing of pain, respectively. However, a model which comprehensively describes how evoked potentials at different latencies – which are the electrophysiological gold-standard for assessing the cerebral processing of pain – and oscillations at different frequencies signal sensory information, expectations, and prediction errors in the processing of pain is lacking so far.

Here, we therefore used EEG to systematically assess the role of evoked potentials and oscillations in the signaling of sensory information, expectations, and prediction errors in the processing of pain. We specifically hypothesized that alpha/beta and gamma oscillations signal predictions and prediction errors, respectively. We further expected that already the earliest evoked responses to noxious stimuli are shaped by predictions, whereas later responses are also shaped by prediction errors. In addition, we speculated that predictions not only shape brain responses to noxious stimuli but also brain activity before a noxious stimulus. To test these hypotheses, we built upon (Geuter et al., 2017), applied a probabilistic cueing paradigm in healthy human participants and performed both frequentist analyses and Bayesian model comparisons.

## Results

To investigate how EEG responses to brief painful stimuli signal stimulus intensity, expectations, and prediction errors (PEs) in the processing of pain, we employed a probabilistic cueing paradigm in 48 healthy human participants. We applied brief painful heat stimuli to the left hand and independently modulated stimulus intensity and expectations in a 2×2 factorial design. To modulate stimulus intensity, we applied painful stimuli of two different levels (high intensity [hi], low intensity [li]). To modulate expectations, the painful stimuli were preceded by one out of two visual cues. The high expectation (HE) cue was followed by a high intensity stimulus in 75% of the trials and by a low intensity stimulus in 25% of the trials. Vice versa, the low expectation (LE) cue was followed by a high intensity stimulus in 25% of the trials and by a low intensity stimulus in 75% of the trials. The experiment, thus, comprised four trial types (Fig. 1a): high intensity, high expectation (hiHE); high intensity, low expectation (hiLE); low intensity, high expectation (liHE); low intensity, low expectation (liLE). In each trial, the participants were asked to provide a rating of the perceived pain intensity on a numerical rating scale ranging from 0 (no pain) to 100 (maximum tolerable pain). In addition, skin conductance responses (SCR) were recorded. Fig. 1b shows the sequence of a single trial.

**Figure 1.**
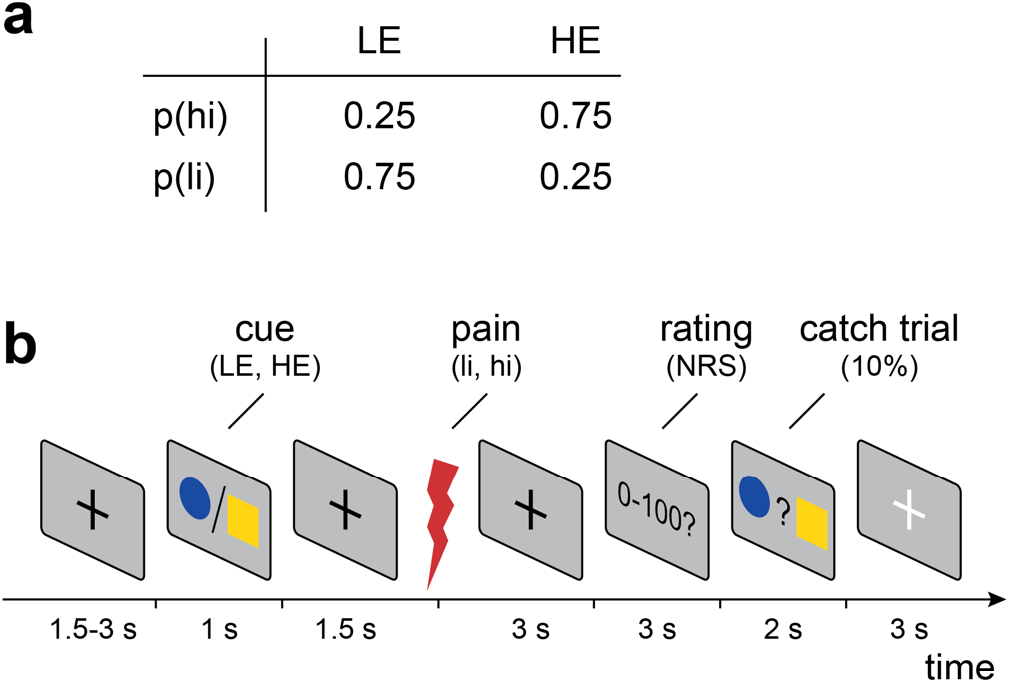
Experimental design. (a) Probabilities of high and low intensity stimuli (p(hi) and p(li), respectively) in high expectation (HE) and low expectation (LE) trials. (b) Each trial started with a central fixation cross with a varying duration of 1.5 to 3 s followed by either a blue dot or yellow square as visual cue. Cues were presented for 1 s and indicated the probability of a subsequent high intensity painful stimulus (0.75 for HE cue or 0.25 for LE cue). The association between the blue dot/yellow square and high intensity (hi)/low intensity (li) painful stimuli was balanced across participants. Next, a painful heat stimulus was applied (3.5 J for high intensity and 3.0 J for low intensity stimuli). Three seconds after the onset of the painful heat stimulus, participants were asked to verbally rate the perceived pain intensity on a numerical rating scale (NRS) ranging from 0 (no pain) to 100 (maximum tolerable pain). In 10% of the trials, a match-to-sample task ensured attention to the cues. In these catch trails, participants were prompted to select the cue that had been displayed during the current trial by a button press. Trials were separated by a break of 3 s during which a white fixation cross was presented.

During the experiment, we recorded EEG and assessed the most consistently observed EEG responses to painful stimuli (Ploner and May, 2018). Evoked EEG responses included the N1, N2, and P2 components. Oscillatory responses included stimulus-induced changes of alpha, beta, and gamma oscillations. In addition, we quantified brain activity before the painful stimulus including the stimulus preceding negativity (SPN; Brown et al., 2008) and oscillatory activity at alpha and beta frequencies.

Building upon previous investigations (Egner et al., 2010; Geuter et al., 2017), we made specific predictions how EEG responses signaling stimulus intensity, expectations, PEs or combinations thereof are modulated across the four trial types (Fig. 2). To formally test these predictions, we pursued two complementary approaches (Egner et al., 2010; Geuter et al., 2017). First, we performed repeated measures analyses of variance (rmANOVAs) with the independent variables stimulus intensity and expectation. In these rmANOVAs, responses signaling stimulus intensity and expectations would manifest as main effects whereas responses signaling PEs would manifest as interactions. To quantify effects and to facilitate interpretation of negative findings, we primarily performed Bayesian rmANOVAs (Keysers et al., 2020). In addition, we performed traditional frequentist rmANOVAs. Detailed results of both Bayesian and frequentist rmANOVAs are provided in Tables 1–3. Second, we employed Bayesian model comparisons based on single-trial data to formally test which combination of stimulus intensity, expectations, and PEs best explained the observed EEG responses. Building upon previous studies (Egner et al., 2010; Geuter et al., 2017), we specifically compared models where stimulus intensity only (INT model), stimulus intensity and expectations (INT+EXP), and expectations and PE (EXP+PE) shaped the respective responses. In line with Geuter et al. (2017), the PE was defined as aversive PE meaning that a prediction error occurs only if the stimulus is more painful than expected. This model has been shown to outperform models with absolute and signed PE formulations (Geuter et al., 2017).

**Figure 2.**
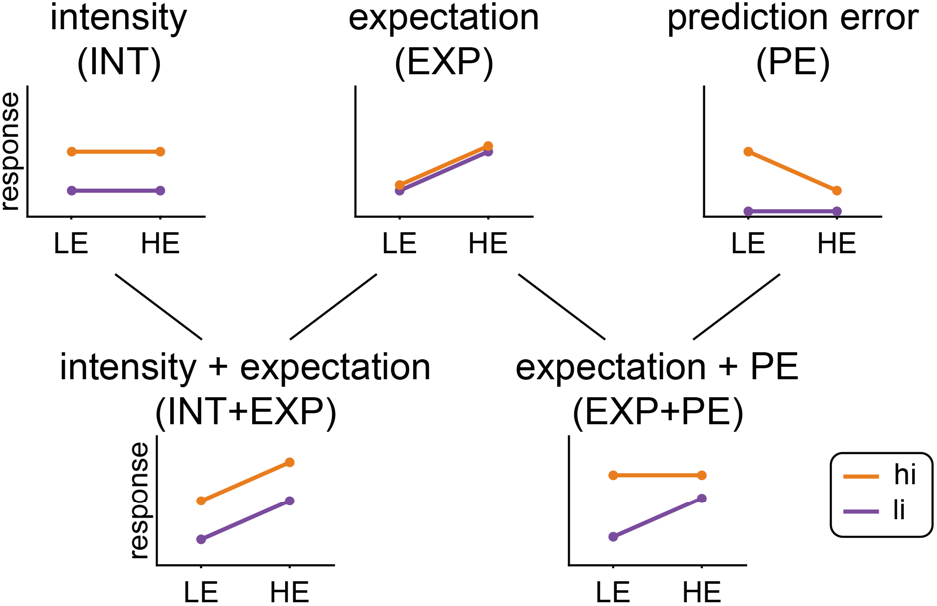
Predicted response patterns for responses signaling stimulus intensity (INT model), expectations (EXP model), prediction errors (PE model) or combinations thereof (INT+EXP model, EXP+PE model).

**Table 1.** Effects of stimulus intensity, expectations and prediction errors on pain ratings and skin conductance responses (SCR). Results of Bayesian and frequentist repeated measures ANOVAs with pain rating and skin conductance response as dependent variables.

| | IV | F | p | $\eta^2$ | BF | mean ± SD | |
| --- | --- | --- | --- | --- | --- | --- | --- |
| pain | intensity | 86.0 | < 0.001 | 0.55 | $1.6e^{26}$ | hiLE | $28.7 \pm 16.6$ |
| | cue | 57.0 | < 0.001 | 0.068 | $1.2e^4$ | hiHE | $35.8 \pm 19.5$ |
| | intensity x cue | 12.8 | < 0.001 | $4e^{-3}$ | 0.36 | liLE | $14.0 \pm 8.9$ |
| | IV df, error df | 1, 47 | | | | liHE | $18.3 \pm 10.5$ |
| SCR | intensity | 51.1 | < 0.001 | 0.53 | $5.0e^{13}$ | hiLE | $0.32 \pm 0.24$ |
| | cue | 6.5 | 0.016 | 0.014 | 0.68 | hiHE | $0.35 \pm 0.23$ |
| | intensity x cue | 0.081 | 0.78 | $2.2e^{-4}$ | 0.24 | liLE | $0.12 \pm 0.11$ |
| | IV df, error df | 1, 30 | | | | liHE | $0.15 \pm 0.13$ |

IV = independent variable; BF = Bayes Factor.

### The effects of stimulus intensity, expectations, and prediction errors on pain ratings and SCR

Before analyzing EEG responses, we investigated the effects of stimulus intensity, expectations, and prediction errors on pain intensity ratings. We therefore calculated rmANOVAs with the independent variables stimulus intensity and expectation. Results are shown in Table 1 and Fig. 3. Bayes factors indicated strong evidence for main effects of stimulus intensity (BF = 1.6e^26^) and expectations (BF = 1.2e^4^). Specifically, pain intensity was higher for high intensity than for low intensity stimuli and higher for high expectation than for low expectation trials. Bayesian rmANOVA showed weak evidence against an interaction of stimulus intensity and expectation (BF = 0.36). To further investigate the relationship between stimulus intensity, expectations, prediction errors, and pain ratings, we tested INT, INT+EXP, and EXP+PE models against each other. The comparisons showed strong evidence that the INT+EXP model explained the data better than the INT (BF = 7.8e^20^) or the EXP+PE (BF = 2.8e^−45^) model. Thus, we found that stimulus intensity and expectations, but not prediction errors shaped pain ratings.

**Figure 3.**
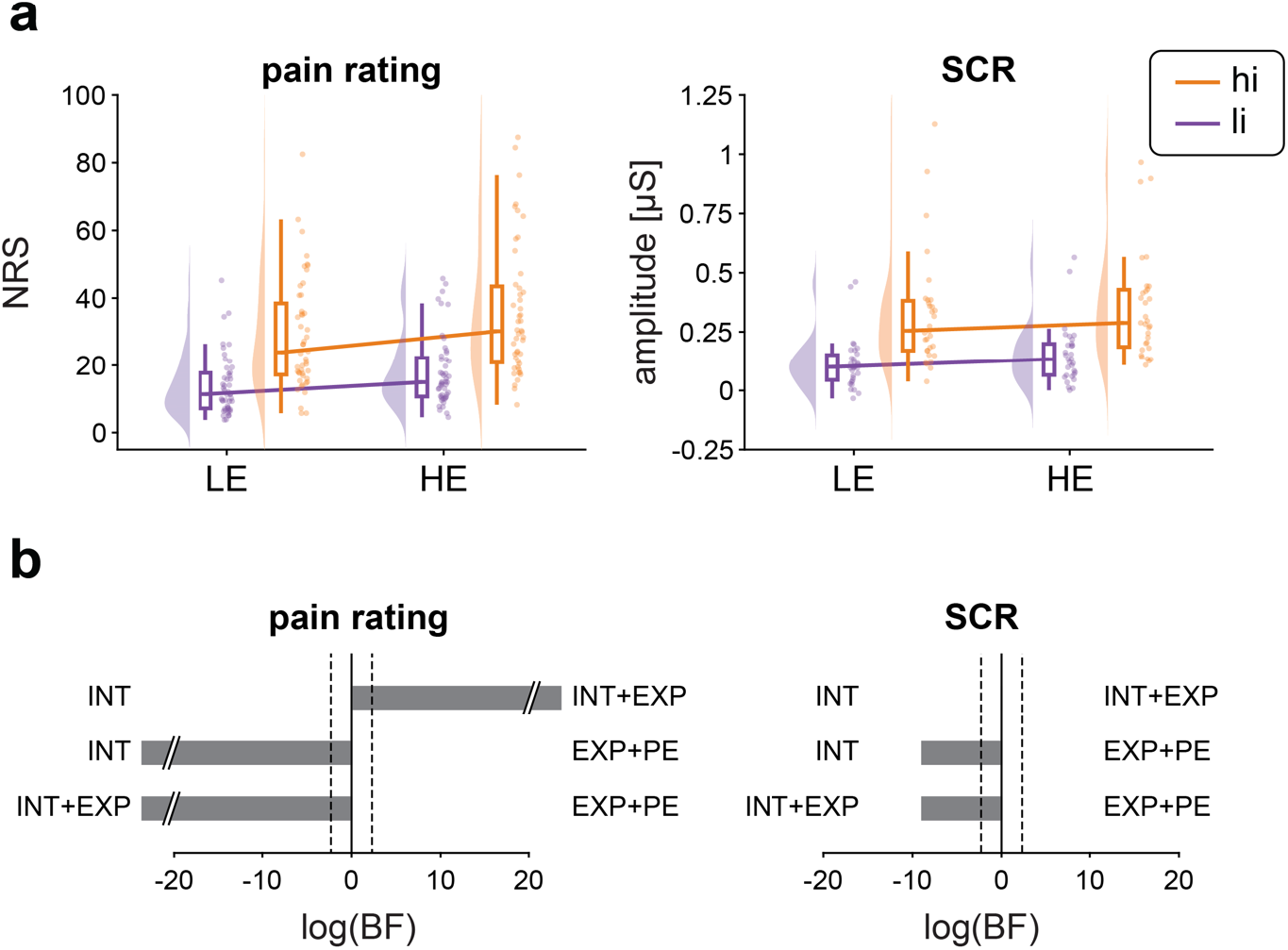
Effects of stimulus intensity, expectations and prediction errors on pain ratings and skin conductance responses (SCR). (a) Raincloud plots (Allen et al., 2019) of pain ratings and SCRs in hiLE, hiHE, liLE, and liHE conditions. The clouds display the probability density function of the individual means, indicated by dots. Boxplots depict the sample median as well as first (Q1) and third quartiles (Q3). Whiskers extend from Q1 to the smallest value within Q1 – 1.5 x interquartile range (IQR) and from Q3 to the largest values within Q3 + 1.5 x IQR. (b) Bayesian model comparisons between stimulus intensity (INT) and stimulus intensity + expectations (INT+EXP) models, stimulus intensity (INT) and expectations + prediction error (EXP+PE) models, and stimulus intensity + expectations (INT+EXP) and expectations + prediction error (EXP+PE) models. Bars depict the natural logarithm of the Bayes factors (BFs). Discontinuous bars indicate log(BF) > 20 or log(BF) < −20. Dotted lines indicate the bounds of strong evidence (log(BF = 0.1) and log(BF = 10)) (Keysers et al., 2020).

We next investigated how stimulus intensity, expectations and prediction errors shaped SCR. The rmANOVA for the SCR showed strong evidence for a main effect of stimulus intensity (BF = 3.2e^12^), i.e. the amplitude of SCRs was higher in high intensity than in low intensity trials (Table 1 and Fig. S1). However, we found inconclusive evidence regarding a main effect of expectation (BF = 0.68) and weak evidence against an interaction of stimulus intensity and expectation (BF = 0.24) on SCR. Bayesian model comparisons of single-trial SCRs showed evidence that the INT model explained the SCR just as well as the INT+EXP model (BF = 1.0) and better than the EXP+PE (BF = 1.4e^−4^) model (Fig. 3).

Taken together, we found strong effects of stimulus intensity on pain intensity ratings and SCRs. Moreover, we found a strong effect of expectations on pain intensity but only inconclusive evidence for an effect of expectations on SCRs. Furthermore, we did not observe an interaction between stimulus intensity and expectation in shaping pain ratings and SCRs.

### The effects of stimulus intensity, expectations, and prediction errors on EEG responses to noxious stimuli

To investigate the effects of stimulus intensity, expectations, and PEs on EEG responses, we calculated rmANOVAs as done for pain intensity ratings and SCR. Bayesian rmANOVAs showed strong evidence for a main effect of stimulus intensity on all EEG responses (BF > 1.2e^3^). N1, N2, and P2 responses (Table 2, Fig. 4) as well as post-stimulus gamma oscillations and alpha- and beta-suppressions (Table 3, Fig. 5) were stronger in the high intensity than in the low intensity conditions.

**Table 2.** Effects of stimulus intensity, expectations and prediction errors on evoked EEG responses to noxious stimuli. Results of Bayesian and frequentist repeated measures ANOVAs with amplitudes of N1, N2 and P2 responses as dependent variables.

| | IV | F | p | $\eta^2$ | BF | mean ± SD | |
| --- | --- | --- | --- | --- | --- | --- | --- |
| N1 | intensity | 30.5 | < 0.001 | 0.36 | 3.4e<sup>11</sup> | hiLE | -4.1 ± 4.6 |
| | cue | 6.0 | 0.018 | 9e<sup>-3</sup> | 0.39 | hiHE | -4.7 ± 4.9 |
| | intensity x cue | 0.92 | 0.34 | 1e<sup>-3</sup> | 0.24 | liLE | -1.2 ± 1.8 |
| | IV df, error df | 1, 44 | | | | liHE | -1.6 ± 2.2 |
| N2 | intensity | 72.0 | < 0.001 | 0.57 | 3.6e<sup>24</sup> | hiLE | -7.7 ± 6.3 |
| | cue | 0.29 | 0.59 | 2e<sup>-4</sup> | 0.17 | hiHE | -7.9 ± 6.5 |
| | intensity x cue | 0.068 | 0.80 | 3.8e<sup>-5</sup> | 0.20 | liLE | -2.0 ± 2.5 |
| | IV df, error df | 1, 47 | | | | liHE | -2.0 ± 2.7 |
| P2 | intensity | 93.8 | < 0.001 | 0.54 | 1.5e<sup>22</sup> | hiLE | 5.1 ± 3.4 |
| | cue | 0.098 | 0.76 | 2.4e<sup>-4</sup> | 0.16 | hiHE | 5.1 ± 3.9 |
| | intensity x cue | 8.1e<sup>-6</sup> | 0.99 | 1.4e<sup>-9</sup> | 0.20 | liLE | 1.8 ± 2.3 |
| | IV df, error df | 1, 47 | | | | liHE | 1.8 ± 2.7 |

IV = independent variable; BF = Bayes Factor.

**Figure 4.**
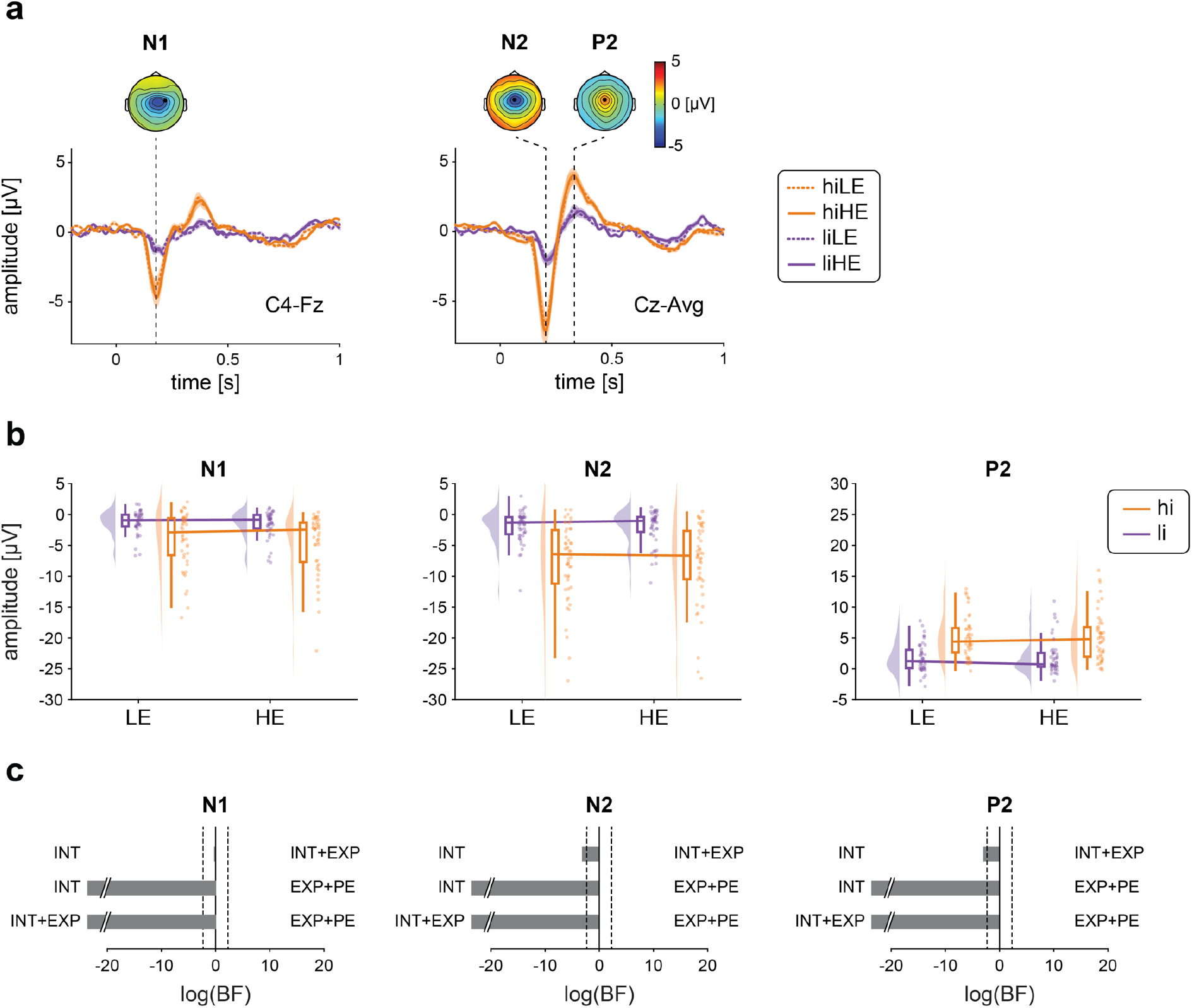
Effects of stimulus intensity, expectations and prediction errors on evoked EEG responses to noxious stimuli. (a) Grand averages of laser-evoked responses. Orange and violet shadings indicate the standard error of the mean. Topographies are based on the average of all four conditions at peak latencies. (b) Raincloud plots (Allen et al., 2019) of N1, N2, and P2 amplitudes in hiLE, hiHE, liLE and liHE conditions. The clouds display the probability density function of the individual means indicated by dots. Boxplots depict the sample median as well as first (Q1) and third quartiles (Q3). Whiskers extend from Q1 to the smallest value within Q1 – 1.5 x interquartile range (IQR) and from Q3 to the largest values within Q3 + 1.5 x IQR. (c) Bayesian model comparisons between stimulus intensity (INT) and stimulus intensity + expectations (INT+EXP) models, stimulus intensity (INT) and expectations + prediction error (EXP+PE) models, and stimulus intensity + expectations (INT+EXP) and expectations + prediction error (EXP+PE) models. Bars depict the natural logarithm of the Bayes factors (BFs). Discontinuous bars indicate log(BF) > 20 or log(BF) < −20. Dotted lines indicate the bounds of strong evidence (log(BF = 0.1) and log(BF = 10)) (Keysers et al., 2020).

**Table 3.** Effects of stimulus intensity, expectations and prediction errors on induced oscillatory EEG responses to noxious stimuli. Results of Bayesian and frequentist repeated measures ANOVAs with amplitudes of alpha, beta and gamma oscillations as dependent variables.

| | IV | F | p | $\eta^2$ | BF | mean ± SD | |
| --- | --- | --- | --- | --- | --- | --- | --- |
| alpha | intensity | 25.4 | < 0.001 | 0.22 | 2.3e<sup>6</sup> | hiLE | 1.2 ± 0.88 |
| | cue | 0.63 | 0.43 | 2e<sup>-3</sup> | 0.22 | hiHE | 1.1 ± 0.80 |
| | intensity x cue | 0.008 | 0.93 | 2.5e<sup>-5</sup> | 0.22 | liLE | 1.5 ± 1.3 |
| | IV df, error df | 1, 47 | | | | liHE | 1.5 ± 1.1 |
| beta | intensity | 11.8 | 0.001 | 0.12 | 1.2e<sup>3</sup> | hiLE | 0.27 ± 0.14 |
| | cue | 0.31 | 0.58 | 2e<sup>-3</sup> | 0.18 | hiHE | 0.27 ± 0.13 |
| | intensity x cue | 0.62 | 0.43 | 2e<sup>-3</sup> | 0.26 | liLE | 0.30 ± 0.17 |
| | IV df, error df | 1, 47 | | | | liHE | 0.30 ± 0.15 |
| gamma | intensity | 10.3 | 0.002 | 0.14 | 4.6e<sup>3</sup> | hiLE | 0.045 ± 0.02 |
| | cue | 3.7e<sup>-4</sup> | 0.99 | 6.6e<sup>-7</sup> | 0.16 | hiHE | 0.045 ± 0.02 |
| | intensity x cue | 9.7e<sup>-5</sup> | 0.99 | 2.6e<sup>-7</sup> | 0.24 | liLE | 0.040 ± 0.01 |
| | IV df, error df | 1, 47 | | | | liHE | 0.040 ± 0.02 |

IV = independent variable; BF = Bayes Factor.

**Figure 5.**
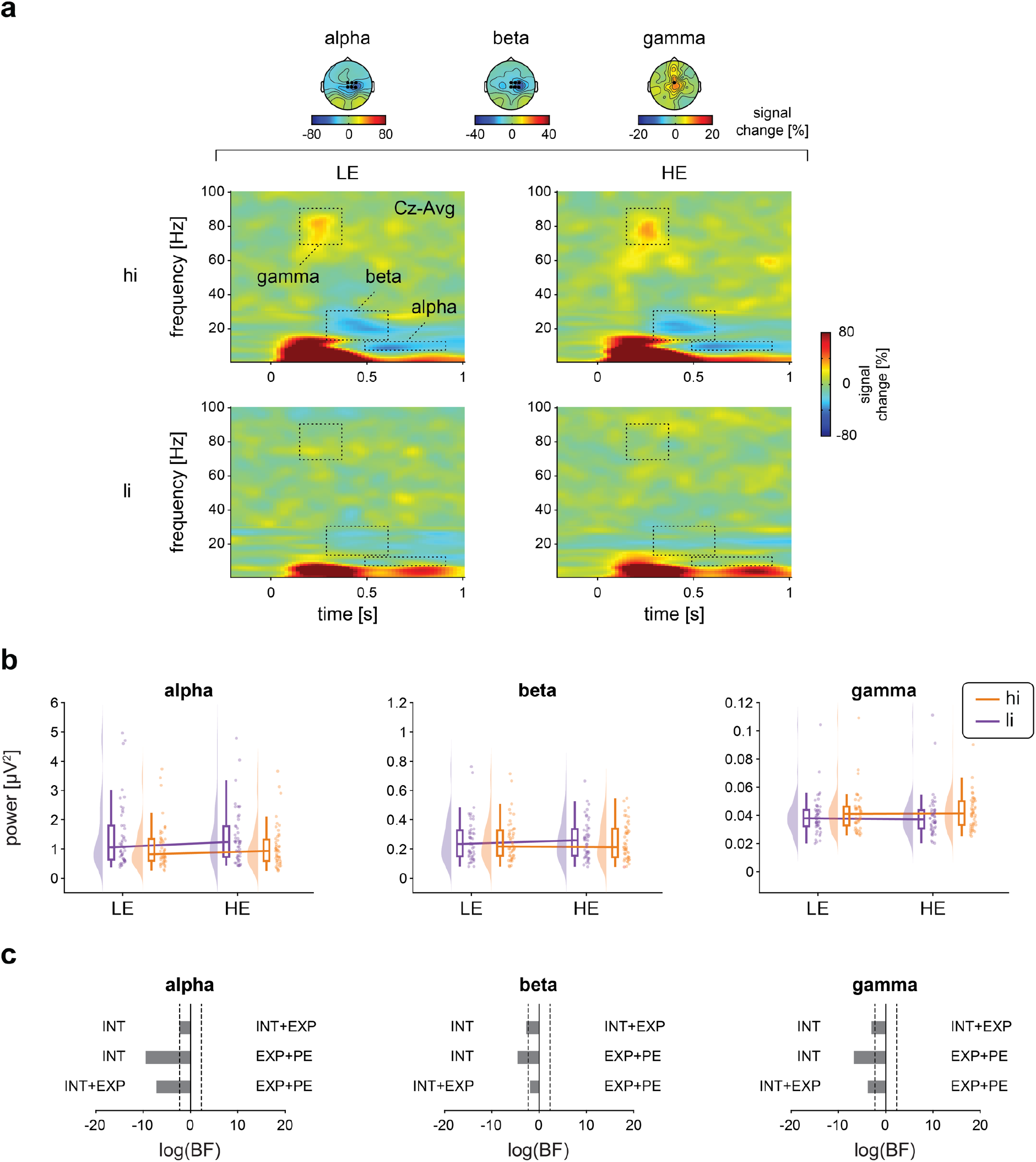
Effects of stimulus intensity, expectations, and prediction errors on oscillatory EEG responses to noxious stimuli. (a) Grand averages of time-frequency representations (TFRs) are depicted as relative change to the baseline preceding the cue presentation (−3.3 to −2.8 s). For visualization TFRs at Cz are presented. Statistical analysis was performed on absolute power values without baseline correction averaged across regions-of-interest (ROIs) as indicated by the dotted boxes and marked electrodes. Topographies display the average of ROIs across all four conditions. (b) Raincloud plots (Allen et al., 2019) of alpha, beta and gamma power in hiLE, hiHE, liLE and liHE conditions. The clouds display the probability density function of the individual means indicated by dots. Boxplots depict the sample median as well as first (Q1) and third quartiles (Q3). Whiskers extend from Q1 to the smallest value within Q1 – 1.5 x interquartile range (IQR) and from Q3 to the largest values within Q3 + 1.5 x IQR. (c) Bayesian model comparisons between stimulus intensity (INT) and stimulus intensity + expectations (INT+EXP) models, stimulus intensity (INT) and expectations + prediction error (EXP+PE) models, and stimulus intensity + expectations (INT+EXP) and expectations + prediction error (EXP+PE) models. Bars depict the natural logarithm of the Bayes factors (BFs). Discontinuous bars indicate log(BF) > 20 or log(BF) < −20. Dotted lines indicate the bounds of strong evidence (log(BF = 0.1) and log(BF = 10)) (Keysers et al., 2020).

In contrast, we found moderate evidence against an expectation effect on all EEG responses (all BF < 0.22) apart from the N1 where evidence was inconclusive (BF = 0.39). In addition, we found moderate evidence against an interaction of stimulus intensity and expectation for all EEG responses (all BF < 0.26). We, thus, observed that the most consistently observed evoked and oscillatory EEG responses to noxious stimuli were shaped by stimulus intensity but not by expectations or prediction errors.

To test for effects on brain activity other than the predefined EEG responses, ANOVAs and cluster-based permutation tests were performed across the post-stimulus time period from 0 s to 1 s, all frequencies, and all channels which corroborated the results at alpha, beta, and gamma frequencies (Fig. S2). Accordingly, model comparisons for N1, N2 and P2, alpha, beta and gamma responses yielded stronger evidence for the INT model than for the INT+EXP (BF < 0.10) and EXP+PE (BF < 0.012) models except for the N1 which showed inconclusive evidence regarding the comparison of the INT and the INT+EXP models (BF = 0.84). Thus, post-stimulus EEG responses were consistently modulated by stimulus intensity but not by expectations.

Having found no expectation effects on EEG responses, we further asked whether the expectation effect on pain ratings can be explained by a *pattern* of the different EEG responses rather than *each* response in isolation. We therefore performed a multiple regression analysis to test whether difference values (HE - LE) of N1, N2, P2, and alpha-, beta-, and gamma-responses together capture the expectation effect on pain ratings. However, the multiple regression model did not significantly explain any variance in the data (F_(6, 44)_ = 0.66, p = 0.68, R^2^ = 0.094).

In summary, results from rmANOVAs and model comparisons convergingly showed that stimulus intensity shapes all EEG responses. In contrast, we found evidence against an effect of expectations and/or prediction errors on EEG responses. Moreover, expectation effects on pain ratings were neither captured by any single EEG response nor by their combination.

### The effects of expectations on EEG activity before the noxious stimuli

Lastly, expectations might not only influence post-stimulus responses to a painful stimulus but also shape brain activity in anticipation of the painful stimulus. We utilized the high temporal resolution of EEG to disentangle these effects. Specifically, we analyzed the SPN reflected by the average amplitude at Cz during 500 ms directly preceding the laser stimulus. In addition, we analyzed oscillatory brain activity during two pre-stimulus phases, i.e. during cue presentation and between cue presentation and painful stimulus (Table 4 and Fig. 6a). Bayesian dependent samples t-tests showed evidence against an expectation effect on the SPN (BF = 0.17). In contrast, expectations significantly influenced oscillatory brain activity at alpha and beta frequencies (Table 4 and Fig. 6b). In particular, the cue-induced decrease in alpha oscillations was stronger for high expectation trials than for low expectation trials (BF = 98.8 for the cue presentation phase, BF 2.0 for the phase between cue presentation and painful stimuli). In addition, the cue-induced decrease in beta oscillations was stronger for high expectation than for low expectation trials (BF = 8.5 for the phase between cue presentation and pain stimulus). These results were corroborated by cluster-based permutation tests which were performed less restrictively across time, frequencies, and all channels (Fig. S2). The cluster-based permutation tests were performed separately for the period of cue presentation and for the period closely preceding the painful stimulus. We, thus, found that expectations shaped neuronal oscillations at alpha and beta frequencies before the painful stimulus.

**Table 4.**
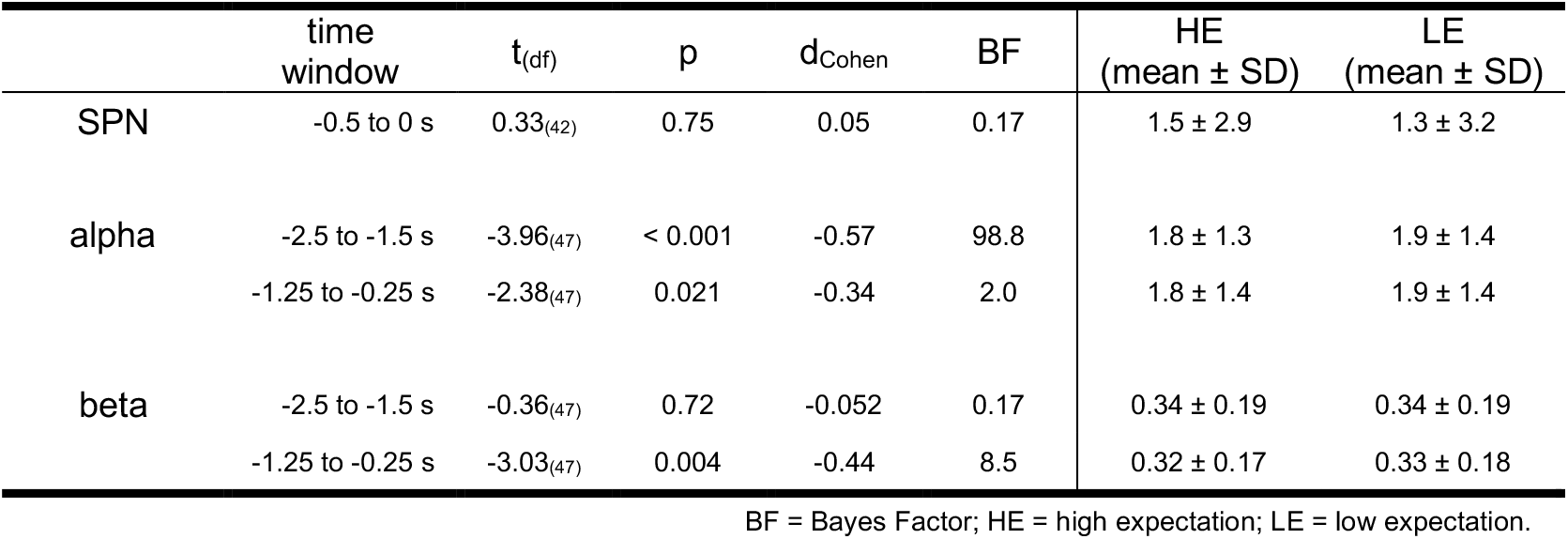
The effects of expectations on EEG activity before the noxious stimuli. Dependent samples t-tests of pre-stimulus brain activity between high expectation and low expectation trials.

**Figure 6.**
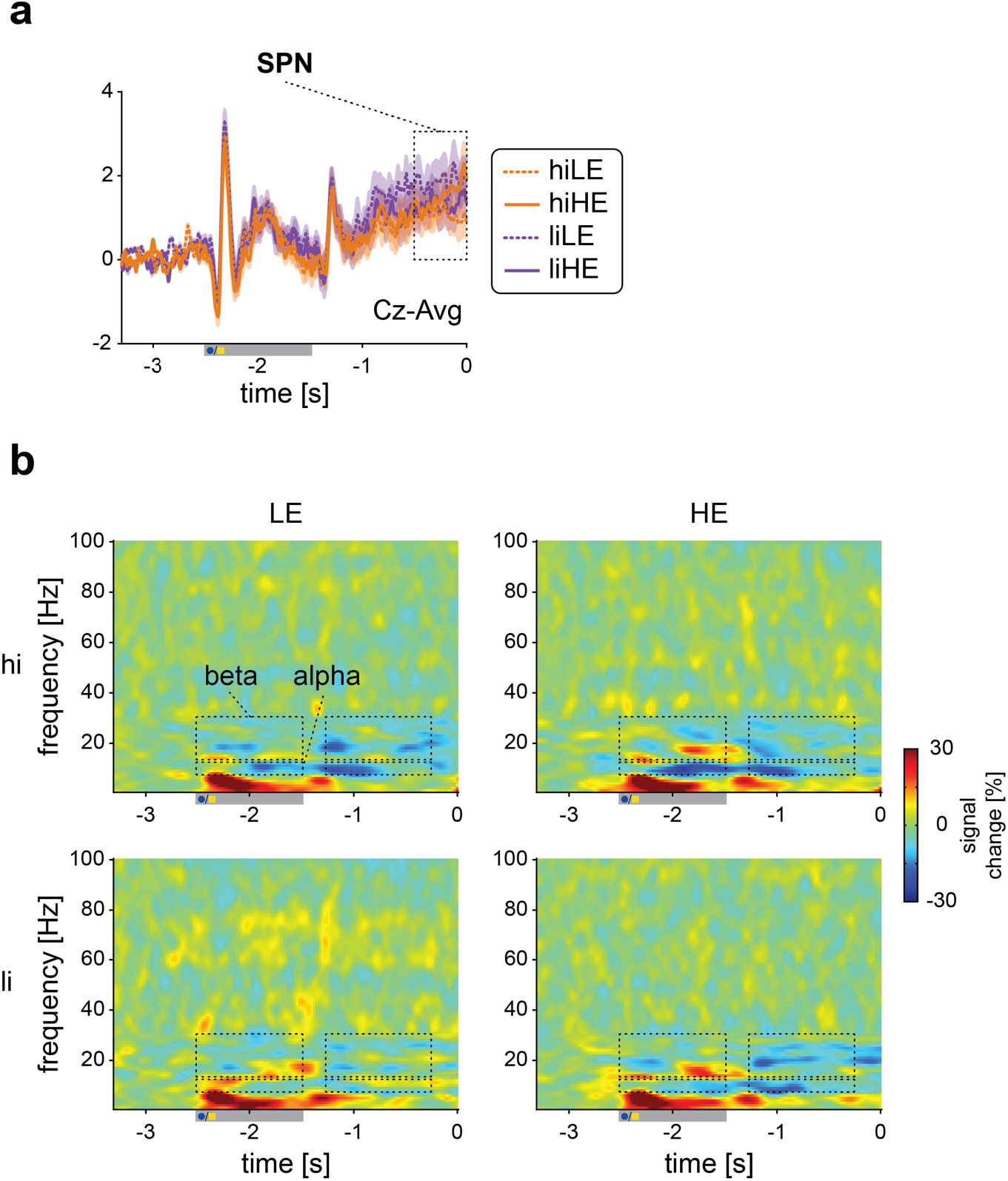
The effects of expectations on EEG activity before the noxious stimuli. (a) Grand averages of time-domain signals at Cz are depicted which preceded the laser stimulus. Orange and violet shadings denote the standard error of the mean. The amplitude of the SPN was determined by averaging the signal at Cz between −0.5 and 0 as indicated by the dotted box. Painful stimuli onset occurs at t=0. The grey bar below the x-axis indicates the time period of cue presentation. (b) Grand averages of the time-frequency representation are depicted as relative change to the baseline preceding the cue presentation (−3.3 to −2.8 s). Dotted boxes represent the two different time windows considered at the alpha and beta frequencies. Painful stimuli onset occurs at t=0. The grey bar below the x-axis indicates the time period of cue presentation. Data were averaged across channels Cz, C2, C4, CPz, CP2, and CP4 and across time and frequency points as indicated by the dotted boxes. Statistical analysis was performed on absolute power values without baseline correction.

## Discussion

In the present study, we observed that sensory information significantly shaped the perception of pain and EEG responses commonly associated with pain. Expectations, in contrast, modulated the perception of pain but not associated EEG responses. Bayesian hypothesis testing confirmed that the observed lack of expectation effects on EEG responses can indeed be interpreted as an absence of effects. These findings indicate that commonly analyzed EEG responses to painful stimuli are more involved in signaling sensory information than in signaling expectations or mismatches of sensory information and expectations. Moreover, they indicate that the effects of expectation on pain are served by different brain mechanisms than those conveying effects of sensory information on pain and are not well captured by commonly analyzed EEG responses to noxious stimuli. We will discuss the implications of these findings for understanding the functional significance of EEG responses to pain, particularly in the context of predictive coding frameworks of brain function, and for understanding how the brain mediates expectation effects on pain.

### The functional significance of pain-associated EEG responses

Our observation that stimulus intensity shapes EEG responses to noxious stimuli is in accordance with previous studies which nearly uniformly showed such effects. Expectation effects, on the other hand, were limited to pain ratings and not found for EEG responses in the present study. At first glance, this contrasts with previous studies which have shown that expectations significantly modulate EEG responses (Wager et al., 2006; Colloca et al., 2008; Iannetti et al., 2008; Morton et al., 2010; Lyby et al., 2011; Huneke et al., 2013; Tiemann et al., 2015; Hird et al., 2018). However, expectation effects in those studies were weaker and less consistent than stimulus intensity effects. Moreover, limited statistical power (Button et al., 2013) and publication bias (Ioannidis et al., 2014) might have resulted in an over-estimation of expectation effects across the literature. Thus, although expectations can, in principle, shape EEG responses, the present findings indicate that these responses are more sensitive to stimulus intensity effects than to expectations. Whether this fundamental difference generalizes from expectations to other contextual modulations of pain remains to be determined.

### Pain-associated EEG responses in a predictive coding model of brain function

As the interaction of sensory evidence and predictions crucially shapes the perception of pain, predictive coding frameworks of brain function have been increasingly applied to the processing of pain (Buchel et al., 2014; Ploner et al., 2017; Tabor et al., 2017; Ongaro and Kaptchuk, 2019; Seymour, 2019). Based on these considerations, recent functional imaging studies have started to investigate how the brain encodes sensory information, predictions, and prediction errors in the processing of pain (Geuter et al., 2017; Fazeli and Buchel, 2018). The results have revealed a spatial dissociation between brain areas encoding stimulus intensity, predictions, and prediction errors. More precisely, a dissociation was found in the insular cortex where posterior parts signalled sensory information whereas anterior parts additionally signalled predictions and prediction errors. The present study was inspired by these investigations and adapted their paradigm for EEG. We particularly aimed to assess how evoked and oscillatory brain responses at different latencies and frequencies encode sensory information, expectations, and prediction errors. Based on previous anatomical and physiological evidence (Arnal and Giraud, 2012; Bastos et al., 2012; Bastos et al., 2015; Michalareas et al., 2016), we specifically hypothesized that alpha/beta and gamma oscillations signal predictions and prediction errors, respectively. We further expected that already the earliest laser-evoked responses are shaped by predictions (Rauss et al., 2011; Bendixen et al., 2012), whereas later responses are also shaped by prediction errors (Stefanics et al., 2018).

Our observed pre-stimulus effects support the idea that alpha/beta oscillations are indeed involved in signaling predictions also in the context of pain. However, Bayesian hypothesis testing of post-stimulus effects provided evidence against the hypothesis that predictions and prediction errors shape evoked and oscillatory responses to noxious stimuli. This contrasts with the results of a recent EEG study (Strube et al., 2021) which showed that post-stimulus alpha/beta oscillations and gamma oscillations were shaped by predictions and prediction errors, respectively. This difference between the previous and the present study might be due to different durations of the employed noxious stimulation models. The previous study used contact-heat stimuli of a few seconds duration whereas the present study used radiant-heat laser stimuli of a few milliseconds duration. Laser stimuli are a standard tool for research on the brain mechanisms of pain and for the clinical assessment of nociceptive pathways (Plaghki and Mouraux, 2003). They yield a highly synchronized activation of nociceptive afferents resulting in a short and clear-cut pain sensation. These stimuli therefore offer the opportunity to not only detect non-phase-locked oscillatory responses but to also to record phase-locked evoked responses and to determine their role in signaling sensory information, expectations, and predictions errors as previously done in other modalities (Todorovic et al., 2011; Todorovic and de Lange, 2012). However, predictive coding concepts propose that the precision of sensory information and predictions crucially determines their weight in further processing (Buchel et al., 2014). Thus, the brief laser stimuli of the present study might yield sensory information with a high precision and weight which, in turn, might result in a relative down-weighting of predictions. Hence, for very brief and clear-cut stimuli, the influence of sensory information might outweigh the influence of predictions and prediction errors on EEG responses.

In the present study, we for the first time performed direct Bayesian model comparisons to assess the role of EEG responses in the signaling of sensory evidence, expectations and predictions errors in the processing of pain. While our results reveal that evoked and oscillatory EEG responses – which are commonly used to assess brain processes related to pain for research and clinical practice – are more involved in signaling sensory evidence than expectations or prediction errors, it is important to note that these findings do neither argue in favor of nor against predictive coding models of brain function. Our findings should, thus, not discourage the application of predictive coding frameworks to the processing of pain but rather encourage the search for brain features – other than the commonly analyzed EEG responses – that signal predictions and predictions errors in the processing of pain.

### Brain mechanisms of expectation effects on pain

Our observation of expectation effects on the perception of painful stimuli without effects on associated EEG responses support that neither evoked nor oscillatory EEG responses to noxious stimuli represent a reliable correlate of pain (Legrain et al., 2011). This dissociation might also be relevant for the search for brain-based biomarkers of pain (Davis et al., 2020). Instead, our findings indicate that EEG responses rather represent a correlate of sensory processing which is not always sensitive to contextual modulations. Thus, other processes not captured by commonly analyzed EEG responses to noxious stimuli likely contribute to contextual modulations of pain. These processes might include cognitive evaluation, pain affect, decision making and reward processing. Such higher-level processes might be less strictly time-locked to noxious stimuli and might therefore not be captured by commonly analyzed EEG responses. Furthermore, these processes might take place in deeper brain areas such as the striatum, medial temporal lobe areas, and the brainstem which are involved in expectation effects on pain (Grahl et al., 2018; Shih et al., 2019; Henderson et al., 2020; Tu et al., 2020) but are not well captured by EEG. In this way, the current EEG findings complement fMRI studies showing that the influence of contextual factors including expectations and placebo effects on pain are mediated by spatial patterns other than those capturing sensory processing (Woo et al., 2015; Woo et al., 2017; Zunhammer et al., 2018). This is also in accordance with previous fMRI studies on predictive coding in the processing of pain which showed that the nociceptive sensitive neurologic pain signature (NPS) was mostly shaped by stimulus intensity rather than expectations (Geuter et al., 2017; Fazeli and Buchel, 2018). Moreover, expectation effects on pain are likely not homogenous. For instance, it has been shown that expectation effects induced by social information and associated learning (Koban et al., 2019) as well as positive and negative expectation effects (Shih et al., 2019) differ fundamentally.

### Conclusions

The present results indicate that commonly analyzed EEG responses to noxious stimuli are more sensitive to sensory processes than to expectations or mismatches between sensory processes and expectations. This finding provides novel insights into the functional significance of the complex spatial-temporal-spectral patterns of brain activity associated with pain. Moreover, our observations might motivate and guide further investigations on how the brain signals sensory information, predictions, and prediction errors in the processing of pain. Understanding these processes might also have implications for understanding the brain mechanisms of chronic pain which have been related to abnormally precise predictions (Edwards et al., 2012; Wiech, 2016; Henningsen et al., 2018).

## Materials and Methods

### Participants

This study was performed in healthy human participants who were recruited through advertisements on an online platform of the Technical University of Munich. Prior to any experimental procedures, all participants gave written informed consent. The study protocol was approved by the Ethics Committee of the Medical Faculty of the Technical University of Munich and pre-registered at ClinicalTrials.gov (NCT04296968). The study was conducted in accordance with the latest version of the Declaration of Helsinki and followed recent guidelines for the analysis and sharing of EEG data (Pernet et al., 2020).

Inclusion criteria were age above 18 years and right-handedness. Exclusion criteria were pregnancy, neurological or psychiatric diseases, severe internal diseases including diabetes, skin diseases, current or recurrent pain, regular intake of medication (aside from contraception and thyroidal medication), previous surgeries at the head or spine, metal or electronic implants, and any previous side effects associated with thermal stimulation.

*A priori* sample size calculations using G*Power (Faul et al., 2007) determined a sample size of 36 participants for a repeated measures analysis of variance (rmANOVA) design with one group and 4 measurements (see below for conditions), a power of 0.95, an alpha of 0.05, and medium effect sizes of f = 0.25. This corresponds to an η^2^ (proportion variance explained) of 0.06 (Correll et al., 2020). Overall, 58 healthy human participants (29 females, age: 24.0 ± 4.3 years [mean ± SD]) were recruited. Nine participants were excluded due to either the absence of pain or low pain ratings (< 10 on a numerical rating scale from 0 (no pain) to 100 (maximum tolerable pain)) during the familiarization run (n = 8), excessive startle responses in response to painful stimulation during the training run (n = 1) or technical issues with the response box used during catch trials (n = 1). The final sample comprised 48 participants (all right-handed, 23 females, age: 23.7 ± 3.4 years). Average clinical anxiety and depression scores obtained using the Hospital Anxiety and Depression Scale (HADS) (Zigmond and Snaith, 1983) were below clinically relevant cut-off scores of 8/21 (Bjelland et al., 2002) (anxiety: 3.0 ± 2.1; depression: 0.9 ± 1.1).

### Procedure

To investigate how noxious stimulus intensity, expectations, and prediction errors relate to the cerebral processing of a painful stimulus and the preceding brain activity, the experiment incorporated two noxious heat stimulus intensities (high and low intensity, hi and li) and two visual cues (high and low expectation cue, HE and LE) resulting in four experimental conditions. High expectation cues were followed by high intensity stimuli in 75% of trials (*high intensity & high expectation condition*, hiHE) and low intensity stimuli in 25% of trials (*low intensity & high expectation condition*, liHE). Conversely, low expectation cues were followed by low intensity stimuli in 75% of trials (*low intensity & low expectation condition*, liLE) and by high intensity stimuli in 25% of trials (*high intensity & low expectation condition*, hiLE; Fig. 1a). The sequence of events for each trial is depicted in Fig. 1b. After a fixation period with a duration of 1.5 to 3 s, a visual cue (blue dot or yellow square) was presented for 1 s. 1.5 s after the offset of the cue presentation, a brief painful heat stimulus was applied. Three s after the noxious stimulus, participants were prompted to rate the perceived pain intensity of the preceding painful heat stimulus on a numerical rating scale ranging from 0 (no pain) to 100 (maximum tolerable pain). To ensure sustained attention to the visual cues, a match-to-sample task was incorporated in 10% of the trials. In these catch trials, HE and LE cues were shown simultaneously after the pain rating and participants were asked to identify the cue of the current trial by a button press (left vs. right, according to the position of the cue on the screen). Participants successfully focused on the task during the whole experiment as indicated by an average accuracy of 95.6 ± 0.1% during the match-to-sample task in catch trials. Trials were separated by 3 s during which a white fixation cross was presented. The experiment consisted of four runs with 40 trials each (hiHE [n = 15], hiLE [n = 5], liLE [n = 15], liHE [n = 5]), resulting in total trial numbers of hiHE [n = 60], hiLE [n = 20], liLE [n = 60], liHE [n = 20]. Runs were separated by short breaks of ~3 mins. Contingencies of visual cues and stimulus intensities, i.e. whether a blue dot or a yellow square predicted high/low intensity stimuli and whether the blue dot/yellow square was presented either on the left or right side of the catch trial screen were balanced across participants.

Prior to the main experiment, we applied a sequence of 10 heat stimuli with different intensities to familiarize the participants with the painful stimulation and the intensity rating procedure. Furthermore, participants were explicitly informed about the contingencies between cues and stimulus intensities and participated in a training run with 16 trials using the same experimental setup and contingencies as in the main experiment. The information and the training run were designed to ascertain that all participants were aware of the contingencies and to minimize learning during the main experiment. During the experiment, participants were seated in a comfortable chair and wore protective goggles and headphones playing white noise to cancel out ambient sounds.

### Stimulation

Painful stimuli were applied to the dorsum of the left hand using a neodymium yttrium aluminum perovskite laser (Nd:YAP, Stimul 1340, DEKA M.E.L.A. srl, Calenzano, Italy) with a wavelength of 1340 nm, a pulse duration of 4 ms, and a spot diameter of ~7 mm (Hu and Iannetti, 2019). Stimulus intensity was set to 3.5 J for high intensity stimuli and 3 J for low intensity stimuli (Hu and Iannetti, 2019). To avoid tissue damage and habituation/sensitization, the stimulation site was slightly changed after each stimulus.

### Recordings and preprocessing

EEG data were recorded using actiCAP snap/slim with 64 active sensors placed according to the extended 10-20 system (Easycap, Herrsching, Germany) and BrainAmp MR plus amplifiers (Brain Products, Munich, Germany). All sensors were referenced to FCz and grounded at Fpz. The EEG was sampled at 1000 Hz (0.1 μV resolution) and band-pass filtered between 0.016 and 250 Hz. Impedances were kept below 20 kΩ.

Preprocessing was performed using BrainVision Analyzer software (v2.1.1.327, Brain Products). EEG data were downsampled to 500 Hz after low-pass filtering with a cutoff frequency of 225 Hz. To detect artifacts and to compute independent component (IC) weights, a 1 Hz high-pass filter (fourth-order Butterworth) and a 50 Hz notch filter removing line noise were applied. EEG data of all runs were concatenated. Independent component analysis based on the extended infomax algorithm was applied to the filtered EEG data ranging from −4.2 s to 3.2 s with respect to laser stimulus onset and resulted in 64 ICs. ICs representing eye movements and muscle artifacts were identified (Jung et al., 2000). Subsequently, the identified ICs were subtracted from the unfiltered EEG and data segments of 400 ms centered around data samples with amplitudes exceeding ±100 μV and data jumps exceeding 30 μV were automatically marked for rejection. Remaining artifacts were identified by visual inspection and manually marked for rejection. All electrodes were re-referenced to the average reference. Finally, data were exported to Matlab (vR2019b, Mathworks, Natick, MA) and further analyses were performed using FieldTrip (v20200128 (Oostenveld et al., 2011). We segmented the EEG data into 7 s-epochs ranging from −4 s to 3 s with respect to the laser stimulus onset. All epochs including marked artifacts or trials in which the laser stimulus was not perceived as painful (pain rating = 0) were excluded from further analysis. To match the number of trials between both hi and both li conditions, respectively, the condition with the lowest trial count was identified for every participant and the same number of trials was randomly drawn from the other conditions (maximum number = 20 trials). Further analyses were based on 17.7 ± 1.6 (range: 14 - 20) trials for hiHE/hiLE conditions and 15.3 ± 4.4 (range: 4 - 20) trials for liHE/liLE conditions for each participant.

Skin conductance data were recorded using two Ag/AgCl electrodes attached to the palmar distal phalanges of the left index and middle finger. Data were recorded using the GSR-MR module with constant voltage of 0.5 V and a BrainAmp ExG MR amplifier (Brain Products, Munich, Germany) with low-pass filtering at 250 Hz and a sampling frequency of 1000 Hz. Subsequent offline analysis included low-pass filtering at 1 Hz using a fourth-order Butterworth filter, downsampling to 500 Hz and a visual artifact inspection. Finally, data were exported and segmented into 14 s-epochs ranging from −4 s to 10 s with respect to the laser stimulus. Identical epochs as for the EEG analyses were selected. Furthermore, we had to exclude additional epochs of skin conductance data comprising marked artifacts. As a result, further analyses of skin conductance data were based on 17.7 ± 1.8 (range: 14 - 20) and 17.6 ± 1.6 (range: 14 - 20) trials for hiHE and hiLE conditions, respectively, and 15.2 ± 4.7 (range: 4 - 20) and 15.2 ± 4.8 (range: 4 - 20) trials for liHE and liLE conditions, respectively.

### Time-domain analysis of EEG data

To quantify the amplitudes of laser-evoked N1, N2, and P2 responses, EEG data were band-pass filtered between 1 and 30 Hz (fourth-order Butterworth) and a baseline correction was applied using the time interval between −3.3 and −2.8 s before the painful stimulus. The selected baseline interval preceded the visual cue to avoid expectation effects during the baseline period. To investigate the amplitude of the N1, the data were re-referenced to Fz. First, the latencies of all laser-evoked responses were assessed for each participant using the average across all trials and conditions. We used a peak/trough detection procedure within the time windows 120 to 200 ms, 180 to 300 ms and 250 to 500 ms (Hu and Iannetti, 2019) for the N1, N2, and P2, respectively. Second, to obtain the amplitudes of the average evoked responses, trials were averaged separately for each condition. Amplitudes of N1, N2, and P2 were assessed by averaging a 30 ms window centered at respective latencies determined in the previous step. Amplitudes of N1 and N2/P2 were extracted at channel C4 (Hu et al., 2010) and Cz, respectively. Finally, single-trial estimates of N1, N2, and P2 amplitudes were obtained accordingly by averaging single-trial data across the same 30 ms windows centered at the latencies identified in step one. For the N1, three participants were excluded from statistical analyses due to a lack of a response in step one.

Besides laser-evoked post-stimulus responses, we were interested in pre-stimulus differences in brain activity induced by the expectation of high or low stimuli. Hence, we investigated the stimulus preceding negativity (SPN) by averaging the amplitude at Cz across the 500 ms interval directly preceding the laser stimulus (Brown et al., 2008). All HE and LE trials were averaged separately for each participant. A low-pass filter with a cutoff frequency of 30 Hz (fourth-order Butterworth) and a baseline correction using the time interval between −3.3 and −2.8 s were applied. No further high-pass filter was applied to take low frequencies of the SPN into account. As a consequence, 5 participants had to be excluded from this analysis due to sweating artifacts which could be corrected by high-pass filtering when laser-evoked responses were analyzed.

### Time-frequency analysis of EEG data

To quantify the power of laser induced oscillatory responses, data were transformed to the time-frequency domain. To this end, we applied a fourth-order Butterworth high-pass filter of 1 Hz and a band-stop filter of 49 to 51 Hz to dampen line noise. Subsequently, a fast Fourier transformation was applied to Hanning-tapered EEG data with a moving time window of 500 ms length for the frequencies from 1 to 30 Hz and a window of 250 ms length for the frequencies from 31 to 100 Hz. The step size was set to 20 ms. We chose a longer window for lower frequencies to retrieve more accurate power estimates including at least 4 cycles for frequencies above 8 Hz. To obtain average responses at different frequency bands, time-frequency data were averaged across trials separately for each of the 4 conditions. Responses at alpha (8-12 Hz), beta (13-30 Hz) (Pernet et al., 2020), and gamma (70-90 Hz) (Tiemann et al., 2018) frequency bands were quantified using the time windows 500-900 ms, 300-600 ms (Mouraux et al., 2003; Ploner et al., 2006) and 150-350 ms (Tiemann et al., 2018), respectively. Alpha and beta power was estimated at sensors Cz, C2, C4, CPz, CP2, CP4 covering the somatosensory cortex (Mouraux et al., 2003; Ploner et al., 2006; Hu and Iannetti, 2019). Gamma power was retrieved at sensor Cz (Hu and Iannetti, 2019). Average responses at the different frequency bands were assessed by calculating the mean power estimates across the selected frequencies, time windows and channels (region-of-interest, ROI). Consequently, we obtained three power values for each condition, i.e. 12 power values for each participant. Single-trial responses of different frequency bands were quantified by averaging across the same time-frequency-sensor selection as for the average responses for each trial.

In addition to oscillatory post-stimulus responses, we were interested in pre-stimulus differences in oscillatory brain activity induced by the expectation of high intensity or low intensity stimuli. Pre-stimulus alpha (8-12 Hz) and beta (13-30 Hz) power were obtained using the mean power across the time windows −2.5 to −1.5 s as well as −1.25 to −0.25 s and the sensors Cz, C2, C4, CPz, CP2, CP4. We chose these time windows to investigate power differences during cue presentation (−2.5 to −1.5 s) and closely before laser stimulus onset (−1.25 to −0.25 s). Data immediately preceding the laser stimulus were not analyzed to avoid confounding pre-stimulus power estimates with post-stimulus activity due to the 500 ms sliding window.

### Analysis of skin conductance data

To quantify skin conductance responses (SCRs), epochs of skin conductance data were averaged across trials for each condition and participant. Amplitudes were defined as peak amplitudes of the maximal peak within a time window from 1 to 7.8 s post-stimulus following a peak detection procedure (Boucsein, 2012; Tiemann et al., 2018) and a baseline correction using skin conductance data at time point zero of the laser stimulus onset. Prior to the quantification of skin conductance responses (SCRs), we identified non-responders by averaging skin conductance data across all trials and conditions for each participant. If the detected global amplitude was below 0.05 μS participants were defined as non-responders (Boucsein, 2012). Based on these criteria, 17 participants were classified as non-responders and excluded from further analysis of SCR data.

### Statistical analyses

Statistical analyses were performed using the statistical software packages JASP (v0.14.1, (JASP) and R (v3.6, (R Core Team, 2020). Motivated by previous findings (Egner et al., 2010; Geuter et al., 2017), we investigated 5 different response patterns (Fig. 2) which vary with respect to the integrated predictors. Specifically, the predictors stimulus intensity (INT), expectation (EXP), prediction error (PE), a linear combination of stimulus intensity and expectation (EXP+INT) as well as a linear combination of expectation and prediction error (EXP+PE) were investigated. The latter has been termed the predictive coding model (Buchel et al., 2014). To investigate these patterns, we computed repeated-measures analysis of variance rmANOVAs for each post-stimulus response (pain rating, N1, N2, P2, alpha-, beta- and gamma-power, SCR) as dependent variable using stimulus intensity (hi vs. li) and cue (HE vs. LE) as factors. Bayesian rmANOVAs were performed as these allow to specifically evaluate evidence for the null hypothesis of no effect (Keysers et al., 2020). In Bayesian rmANOVAs, the Bayes Factor (BF) is defined as ratio between the likelihood of the data given a model including the effect of interest and the likelihood of the data given an equivalent model with the effect of interest removed. BF < 1/3 and BF < 1/10 indicate moderate and strong evidence in favor of the absence of the effect of interest, respectively. BF > 3 and BF > 10 indicate moderate and strong evidence in favor of the effect of interest, respectively (Keysers et al., 2020). For all Bayesian statistics, default Cauchy priors were chosen in JASP. Complementing Bayesian inference, we also conducted frequentist rmANOVAs with the same factors. Post-hoc dependent samples t-tests were performed if a statistically significant interaction was observed. To test for additional effects outside the predefined ROIs, we performed cluster-based permutation tests across time, frequencies and all channels (see supplementary material for details).

Furthermore, on a single-trial level, we performed formal pairwise model comparisons to test which model explains post-stimulus responses best. Models including stimulus intensity and expectation (INT+EXP) as well as expectation and prediction error (EXP+PE) as predictors were compared to the model solely including stimulus intensity (INT), respectively. Additionally, the EXP+PE model was compared with the INT+EXP model. Linear mixed effects models included either stimulus intensity, expectation, prediction error or combinations thereof as fixed effects and participants as random effects. In contrast to the rmANOVA approach, linear mixed effects models are based on single-trial data and account for differences in trial numbers and variability between subjects. Moreover, the different models can be explicitly formulated and compared. We used the R package BayesFactor (v0.9.12) (Rouder and Morey, 2012) to compute Bayes factors for model comparisons. These Bayes factors quantify the evidence for one model over another model as a ratio of two likelihoods, i.e. the likelihood of the data given each model. Stimulus intensity was coded as 1 for high intensity stimuli and as 0 for low intensity stimuli. Expectation was coded as the probability of a following high intensity stimulus, i.e. 0.75 for high intensity cue conditions (hiHE and liHE) and 0.25 for low intensity cue conditions (hiLE and liLE). Finally, the prediction error was defined as aversive prediction error meaning that a prediction error occurs only if the outcome (stimulus intensity) is more painful than expected. Specifically, the aversive PE was selected because a previous study by Geuter et al. (2017) demonstrated that models incorporating the aversive PE explained brain responses to pain better than absolute and signed PEs. Hence, it was coded as difference between stimulus intensity and expectation, i.e. PE = 1 - 0.25 for hiLE, PE = 1 - 0.75 for hiHE and PE = 0 for for liHE and liLE (Geuter et al., 2017).

To complement univariate analyses using single post-stimulus responses as outcome variables and to investigate whether a combination of N1, N2, P2, alpha-, beta- and gamma-power can predict the expectation effect on pain ratings, we computed difference values of pain ratings, N1, N2, P2, alpha-, beta- and gamma-power by subtracting average values of the LE trials from HE trials for each participant. Subsequently, we tried to predict difference values of pain ratings (dependent variable) based on difference values of N1, N2, P2, alpha-, beta- and gamma-power (independent variables) using multiple regression. Prior to the analysis, all difference values were z-transformed across participants to adjust the data to the same scale.

Finally, we investigated whether cue-induced expectations affected brain activity preceding the laser stimulus. To this end, we performed Bayesian dependent samples t-tests comparing the average amplitude of the SPN between HE and LE trials. Similarly, we compared alpha- and beta-power during two pre-stimulus windows, one during cue presentation and one closely preceding the painful laser stimulus. Again, these tests were accompanied by Bayesian dependent samples t-tests to estimate the evidence for the null-hypothesis. To test for additional effects outside the predefined ROIs, we performed cluster-based permutation tests across time, frequencies and all channels (see supplementary material for details).

## Data availability

All data in EEG-BIDS format (Pernet et al., 2019) and code are available at https://osf.io/jw8rv/.

## Acknowledgments

The study was supported by the Deutsche Forschungsgemeinschaft (PL 321/14-1) and the the European Research Council (ERC) under the European Union’s Horizon 2020 research and innovation programme (grant agreement No 758974).

## Supplementary material

**Figure S1.**
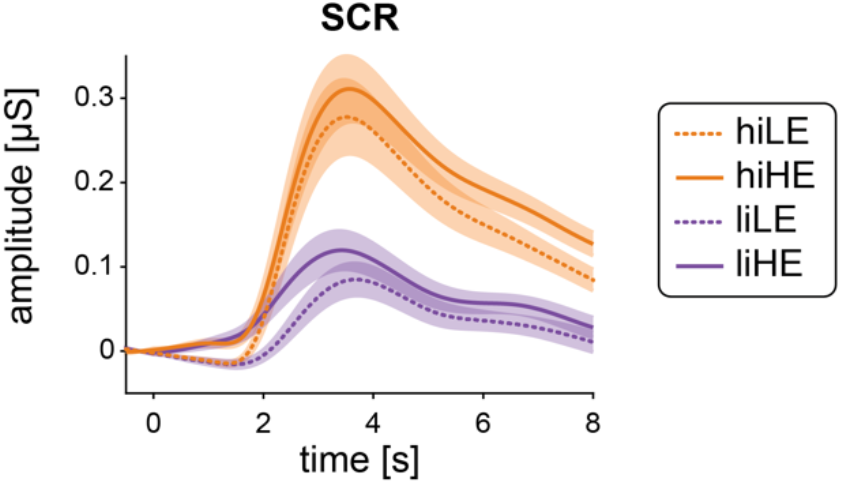
Effects of stimulus intensity, expectations, and prediction errors on skin conductance responses to noxious stimuli. Grand averages of skin conductance responses. Orange and violet shadings indicate the standard error of the mean. HE, high expectation; hi, high intensity; LE, low expectation; li, low intensity; SCR, skin conductance responses.

### Methods – cluster-based permutation tests

Nonparametric cluster-based permutation tests (Maris and Oostenveld, 2007) were performed on three different intervals, two pre-stimulus intervals corresponding to the duration of cue presentation (−2.5 to −1.5 s) and the interval closely before laser stimulus onset (−1.25 to −0.25 s) and one post-stimulus interval from 0 to 1 s after the laser stimulus. Spectral estimates were clustered across time, frequencies, and channels. To test for main effects, all hi, li, LE, and HE trials were pooled separately for each participant and averaged resulting in 4 average time-frequency representations (TFR) per participant. To test for the interaction, we subtracted the liLE from the hiLE (ΔLE) condition and the liHE from the hiHE (ΔHE) condition after averaging across trials within each condition for each participant. Dependent samples t-tests were performed on each time-frequency-channel value. Resulting t-values were thresholded at α = 0.05 and values exceeding the threshold were summed up across neighboring time-frequency-channel values. The minimum number of channels for a cluster was set to 2. To determine the p-value of each cluster, the TFRs hi and li, LE and HE as well as ΔLE and ΔHE were swapped randomly for each participant and the aforementioned statistical procedure was applied. During each of the 1000 permutations, the maximum summed t-value was kept to establish a distribution under the null hypothesis.

### Results – cluster-based permutation tests

#### The effects of stimulus intensity, expectations, and prediction errors on post-stimulus oscillatory brain activity

We observed a main effect of stimulus intensity during the post-stimulus interval as indicated by two significant positive clusters and one significant negative cluster (Fig. S2). The positive cluster at lower frequencies (1 to 18 Hz, p = 0.002) mostly reflected the evoked potential, whereas the positive cluster at higher frequencies (49 to 100 Hz, p = 0.02) represented the gamma response to the painful laser stimuli. Both positive clusters indicate higher power in hi trials than in li trials. In contrast, the significant negative cluster (3 to 36 Hz, p = 002) indicates a stronger decrease in power at alpha and beta frequencies in hi trials as compared to li trials. No additional significant clusters for the main effects of stimulus intensity or expectations (p > 0.39), or the interaction stimulus intensity x expectations (p > 0.71) were found in the post-stimulus interval.

**Figure S2.**
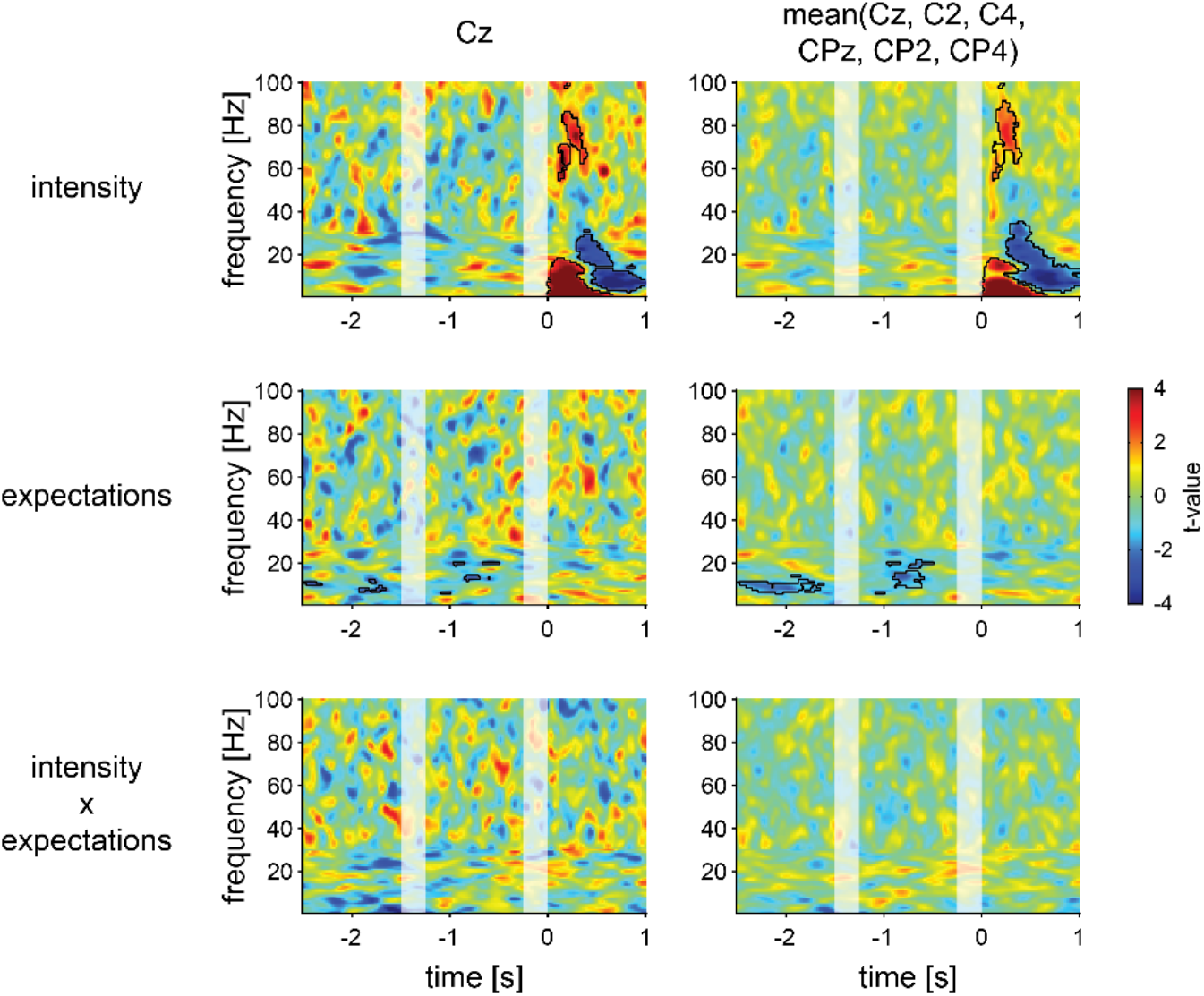
Effects of stimulus intensity, expectations, and prediction errors on pre-stimulus and post-stimulus oscillatory brain activity. T-maps of the cluster-based permutation tests across time, frequencies, and channels at Cz (left column) and averaged across the channels Cz, C2, C4, CPz, CP2, and CP4 (right column). Main effects of stimulus intensity and expectations as well as the interaction stimulus intensity x expectations were tested. Cluster-based permutation tests were performed separately for two pre-stimulus intervals of 1 s each and one post-stimulus interval of 1 s (saturated colors). Significant clusters are outlined in black. Seemingly non-contiguous clusters are connected via additional channels (not depicted). For visualization purposes, time-frequency data points in the right column were marked as part of a significant cluster if the cluster included at least one of the channels averaged.

#### The effects of expectations on pre-stimulus oscillatory brain activity

We observed one significant negative cluster (5 to 15 Hz, p = 0.02) during the cue presentation period which indicated a stronger decrease in alpha power in HE trials as compared to LE trials (Fig. S2). In addition, we found a second significant negative cluster (6 to 22 Hz, p = 0.048) during the period closely preceding the laser stimulus affecting alpha and beta power. Thus, alpha and beta power during the pre-stimulus interval was lower for HE than for LE trials.

